# Endosomal Microautophagy is Activated by Specific Cellular Stresses in Trout Hepatocytes

**DOI:** 10.1101/2024.09.26.615173

**Authors:** Emilio J. Vélez, Vincent Véron, Jeanne Gouis, Steffi Reji, Karine Dias, Amaury Herpin, Florian Beaumatin, Iban Seiliez

**Affiliations:** Université de Pau et des Pays de l’Adour, E2S UPPA, INRAE, UMR1419 Nutrition Métabolisme et Aquaculture, Saint-Pée-sur-Nivelle, France, 64310; INRAE, UR1037 Laboratory of Fish Physiology and Genomics, Rennes, France, 35700

## Abstract

Endosomal microautophagy (eMI) is a recently discovered autophagic process where cytosolic proteins are selectively captured in late endosome/multivesicular bodies (LE/MVB). This pathway, similar to chaperone-mediated autophagy (CMA), involves the recognition of KFERQ-like motif containing proteins by HSC70. While CMA targets substrates to lysosomes via the receptor LAMP2A, eMI involves internalization into intraluminal vesicles within LE/MVB through interactions with ESCRT machinery. Although the same proteins could be targeted by either pathway, eMI’s role in cellular homeostasis is less understood. Our research identified an eMI-like process in rainbow trout hepatocytes, triggered by oxidative stress, high-glucose, DNA damage, and nutrient deprivation, but not serum deprivation. This finding suggests eMI’s stimulus-specific induction and its potential compensatory role when CMA is impaired. Our study provides new insights into eMI and offers novel model organisms for exploring its interactions with CMA, enhancing our understanding of cellular stress responses.

## Introduction

Endosomal microautophagy (eMI) is a recently discovered autophagy process in which cytosolic proteins are selectively trapped in vesicles formed at the membrane level of late endosomes (LE)^1^. eMi can be considered as the “sister” pathway of the more studied chaperone-mediated autophagy (CMA)^2^, as both are characterized by the selective targeting of cytosolic proteins containing a pentapeptide sequence biochemically similar to KFERQ (lysine-phenylalanine-glutamate-arginine-glutamine)^3^. Both processes initiate with the recognition of a KFERQ-like motif by HSC70 (heat shock protein family A (Hsp70) member 8, also known as HSPA8) and co-chaperones^4–6^. For CMA, after binding to the limiting CMA receptor LAMP2A (lysosomal associated membrane protein 2A)^7,8^, the substrates are targeted and translocated into lysosomes for degradation. In contrast, during eMI, the substrate is targeted to the LE membrane via the binding to negatively charged phosphatidylserine residues^2^. The cargo is then internalized thanks to the mediation of some members of the ESCRT machinery (Tsg101 and Alix) and the ATPase Vps4, forming multivesicular bodies (MVB)^1,9,10^. Afterwards, the KFERQ-containing protein can be degraded directly within LE/MVB, or after the fusion of these with lysosomes. Otherwise, the substrate can also be secreted after the fusion of LE/MVB with the plasma membrane^11–13^.

Although the characteristic KFERQ-like motif, required for eMI and CMA targeting, is present in up to 75-80% of the total proteome from yeast to mammals^14^, it is particularly noteworthy that this motif is present in key proteins involved in the control of transcription, cell cycle and cellular energetics, among others key cellular processes^6,14^. Thus, through the selective degradation of those motif-bearing proteins, eMI could play a critical role in maintaining cellular homeostasis and metabolism, as demonstrated for CMA^15–22^. In this regard, CMA malfunction has been associated with diverse pathological situations^23–27^, and similarly, the reduction of eMI during aging has been associated with the progression of some proteinopathies^12,28,29^. Consequently, the interaction between both pathways in targeting KFERQ proteins could be a key step in the adequate remodeling of the cellular proteome during health and disease. For example, the KFERQ-containing protein TAU, linked to cellular senescence and brain disorders (tautopathies)^30^, can be degraded through both CMA and eMI^29^, and during CMA impairment, acetylated TAU is rerouted to eMI^12^. Therefore, exploring the specificities of the regulation of both pathways could reveal novel aspects about the control of metabolism and cellular homeostasis. For instance, eMI and CMA are reciprocally regulated during sustained starvation^9,31^, pointing to a specialization of functions, action scenarios or preferred targets between these two autophagic pathways that deserve further research.

Recently, we provided functional evidence for the existence of a CMA pathway in fish, opening new perspectives for approaching this function from both novel and evolutionary perspectives^27^. This discovery highlighted the compelling need of using new models organisms, such as zebrafish (*Danio rerio*) – a powerful model for functional studies – and rainbow trout (RT, *Oncorhynchus mykiss*) – a model species with impaired glucose tolerance – to better decipher the fundamental processes engaged in this function^20–22^. In contrast, the ability of fish to perform eMI has so far not been reported nor explored^32^. However, our latest research in RT revealed that *in vitro* silencing of the CMA-limiting factor *lamp2a* led to significantly increased levels of the endosomal sorting complexes required for transport-I (ESCRT-I) protein tumor susceptibility 101 (Tsg101)^20^, critical for eMI in mammals^1^. This suggested that an eMI-like process may compensate for impaired CMA activity, as seen in mammals^9,12^. Here, using a fluorescent eMI reporter originally developed for fruit flies^33^, we identified and characterized an eMI-like pathway in a RT hepatocyte cell line. Under mild-oxidative stress, the fluorescent biosensor accumulated in puncta that co-localized with LE. Interestingly, while the formation of these puncta is directly related to the ESCRT machinery, it has noteworthy been shown to be totally independent of CMA and macroautophagy (MA). We also found the RT eMI-like process was triggered by hyperglycemic stress, DNA damage and nutrient deprivation, but not by foetal bovine serum (FBS) removal, showing selective induction. In summary, these results demonstrate for the first time the presence of an eMI-like process in fish, offering new opportunities and models to study this autophagic pathway and its interactions with CMA.

## Results and Discussion

### The core eMI machinery is expressed in fish tissues and RTH-149 cells

eMI process requires KFERQ-like motif recognition by HSC70 and interaction with other key components, including the ESCRT-I protein TSG101 and its associated protein ALG-2 Interacting Protein X (ALIX), the ATPase Vacuolar Protein Sorting 4 (VPS4), as well as the chaperone protein Bag Cochaperone 6 (Bag6)^9^. While HSC70 is highly conserved across phyla^32^, the presence of other eMI components in fish genomes has not been evaluated. Interestingly, during evolution, teleost fish underwent a third round of genome duplication (Ts3R). Additionally, salmonids, including RT, experienced a four round of whole-genome duplication (Ss4R) (Figure 1A)^34^. As a result, up to four orthologs of each human gene might be expected in the RT genome. Homology-based searches in the RT genome (Ensembl release 112) identified several genes with high homology to human *TSG101, VPS4A, VPS4B, ALIX* and *BAG6*. While only one ortholog of *TSG101* has been retained in the RT genome (Figure 1B), we identified two orthologs for *VPS4A* and *ALIX*, and four copies for *VPS4B* and *BAG6* (Figure 1B). Using the *PhyloFish* RNA-seq database, which provides gene expression data from different ray-finned fish species^35^, we screened for expression of those genes across different RT tissues (Figure 1C). We detected mRNA expression of all genes, particularly in the brain, gills, kidney and ovary, with levels varying by tissue and gene. We also analyzed the expression of those genes in the RT liver cell line RTH-149 (Figure 1C red inset), finding that *tsg101, vps4a*C2, *vps4a*C1, *vps4b*C8, *vps4b*C28, *Alix*C18, and the four RT *bag6*s are expressed in this model. Thus, the RTH-149 cell line emerges as an interesting system to further dissect this process using *in vitro* tools. Taken together, these results reveal that the RT displays and express the core eMI machinery, suggesting that this species has the genetic potential to perform an eMI-like process. Moreover, using the same strategy, we have identified corresponding orthologs in zebrafish (Supplementary Figure 1A) and confirmed their tissue-dependent expression (Supplementary Figure 1B). These results suggest that the potential to perform an eMI-like process is not exclusive to RT but a common feature of teleost fishes.

**Figure 1.**
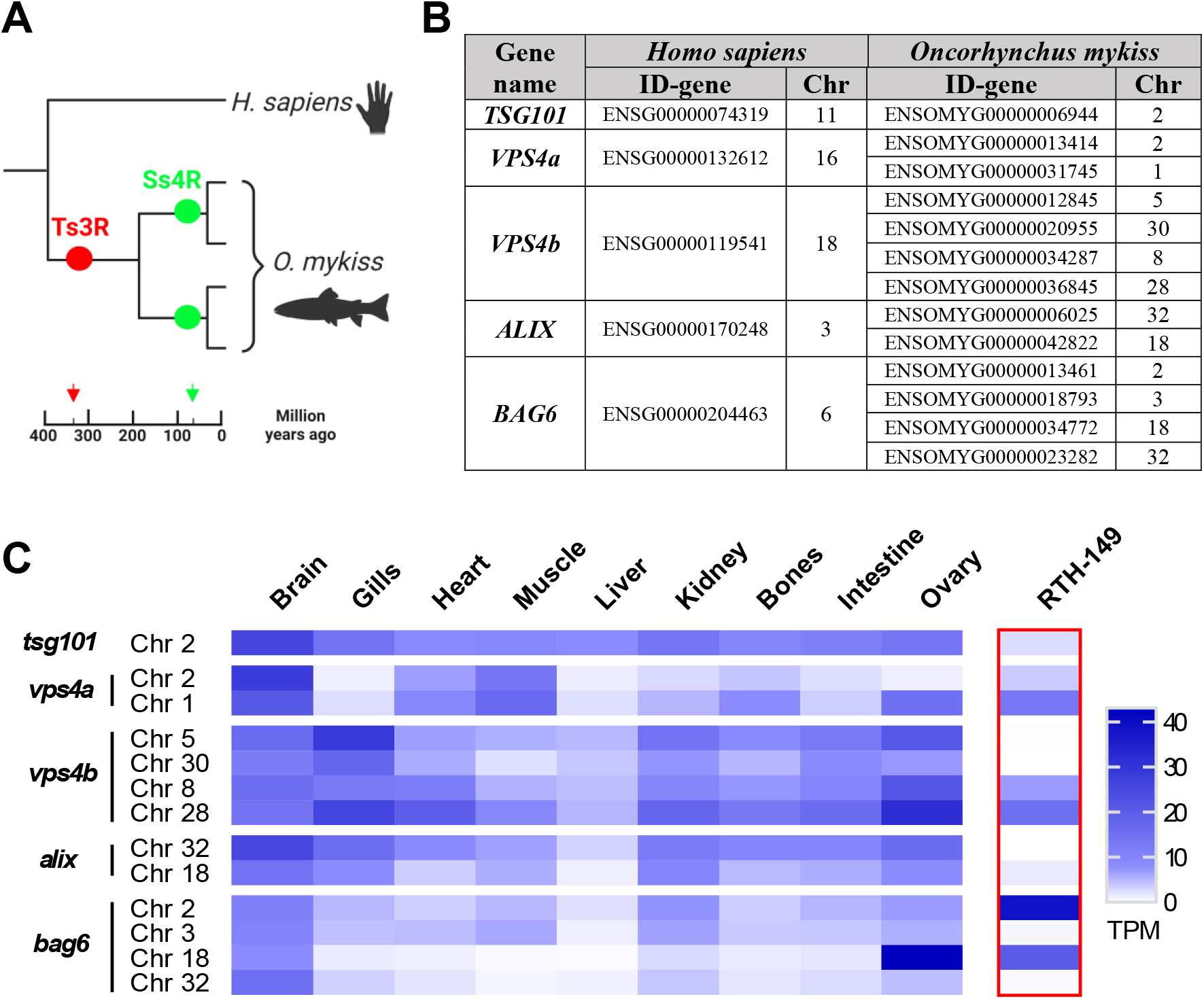
The core eMI machinery is present and expressed in *Oncorhynchus mykiss* (RT, rainbow trout) tissues and the RT-derived RTH-149 cells. (**A**) Schematic representation of the evolutionary history of RT genes showing the third round of genome duplication (Ts3R) experienced by teleost fish, and the additional round of whole-genome duplication (Ss4R) of salmonids. (**B**) Identification (Ensembl ID) and chromosome location (Chr) of the main eMI-genes (*TSG101, VPS4A, VPS4B, ALIX* and *BAG6*) in the human genome, and their corresponding RT orthologs. (**C**) Heat map showing mRNA expression levels (TPM, transcripts per million) of the different identified genes in different tissues of RT (brain, gills, heart, muscle, liver, kidney, bones, intestine and ovary) extracted from the RNA-Seq database PhyloFish, as well as in RTH-149 cells (framed with red line) analyzed by BRB-Seq in the current work.

### Mild-oxidative stress induces eMI-like puncta that co-localize with LE in RTH-149 cells

Recently, we demonstrated that mild-oxidative stress induces recognition of a KFERQ-fluorescent reporter by chaperones for its targeting and degradation by CMA in RTH-149 cells^20^. Here, we have used an eMI-reporter previously employed to track eMI in flies and mammals^29,33^. It consists in a KFERQ recognition motif fused to either the N-terminal or C-terminal of the split Venus. While hemi KFERQ-tagged Venus proteins can be unfolded and rapidly degraded in lysosomes by CMA, resulting in no fluorescence, they would fluoresce only when the N- and C-terminal parts are reconstituted in close proximity within the MVB during eMI^29^. Here, we generated RTH-149 cells that stably express both KFERQ-split Venus proteins, and tested whether mild-oxidative stress (H_2_O_2_, 25 μM) could induce an eMI-like process, as observed in flies^36^. Under control (CT) conditions, the eMI sensor displayed a diffuse distribution with only few discrete and scattered puncta (Figure 2A; quantification in Figure 2B). In contrast, H_2_O_2_ treatment for 8 or 16 h resulted in higher numbers of Venus puncta, showing a clear time-response effect (Figure 2A; quantification in Figure 2B). To locate these H_2_O_2_-induced Venus puncta, we assessed their co-occurrence with the LE marker RAB7 (Ras-related protein 7)^9,37^ labelled with RFP. We found a high level of co-occurrence between eMI-sensor puncta induced by mild-oxidative stress and the RAB7 signal (Figure 2C). The positive Pearson’s correlation coefficient between the intensity of the two fluorescent channels (PCC, Mean ± SEM: 0.914 ± 0.004 from a total of 85 cells RAB7+) supported their co-occurrence^38^.

**Figure 2.**
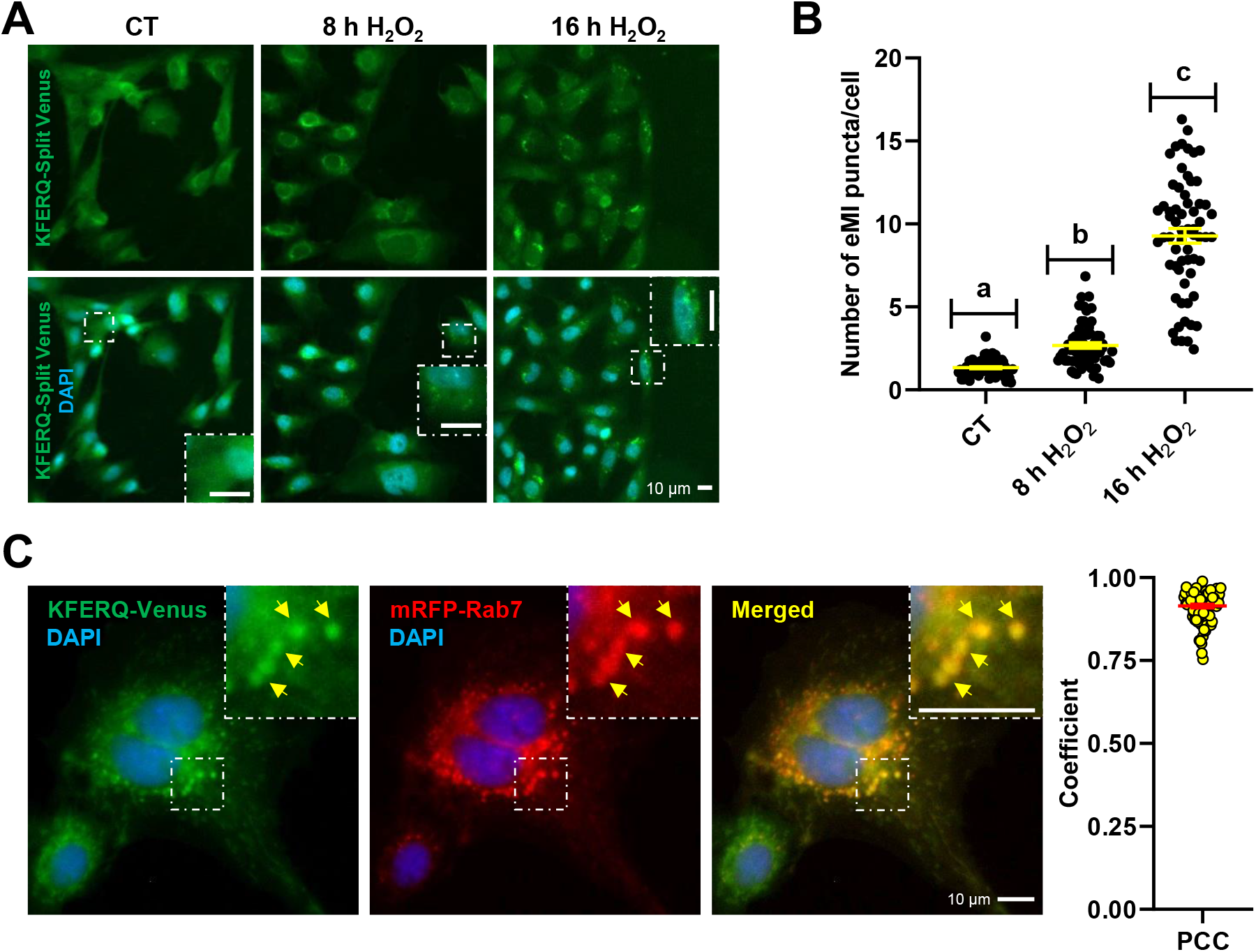
Mild-oxidative stress (H_2_O_2_ 25 μM) induced the targeting of the eMI-reporter Venus to late endosomes through an eMI-like process in RT cells. (**A**) Representative images of RTH-149 cells stably expressing an eMI-reporter consisting in a KFERQ recognition motif fused to either the N-terminal or C-terminal of split Venus and visualized by fluorescence microscopy. The exposure to H_2_O_2_ 25 μM, for either 8 h or 16 h, induced the reconstitution of the Venus fluorescent protein (green) and the formation of Venus-puncta, in comparison with the CT, where the reporter showed a diffuse distribution with only discrete and scattered puncta standing out from the background noise. Nuclei were stained with DAPI (blue) and the inset show an amplified region of each image. (**B**) Quantification of KFERQ-Venus number of puncta per cell. All values correspond to individual images (CT 63; 8 h H_2_O_2_ 63; 16 h H_2_O_2_ 63), with >18 images/experiment in a total of 3 independent experiments (> 600 cells per condition). Different letters denote significant differences between groups compared by the non-parametric Kruskal-Wallis test (p < 0.0001) followed by Dunn’s multiple comparisons tests. All data are presented as Mean ± SEM; scale bars: 10 μm. (**C**) Representative images of H_2_O_2_-induced KFERQ-Venus puncta (green) in cells transiently transfected with RAB7-RFP (red) visualized by fluorescence microscopy. Most of the KFERQ-Venus puncta co-localized with the red late endosome marker RAB7-RFP, as indicated by the yellow arrowheads and amplified in the insets, and as supported by the positive Pearson’s correlation coefficient (PCC) results of the global analysis of 85 single cells using the BIOP version of JACoP plugin for Fiji. Scale bars: 10 μm.

These results suggest that mild-oxidative stress in RT cells induces the targeting of KFERQ-containing proteins to LE in a process similar to eMI in flies and mammals^6^.

### The occurrence of eMI-like puncta under mild-oxidative stress relies on ESCRT machinery but not on CMA nor on MA

To determine if the Venus puncta co-occurring in response to oxidative stress corresponds to an indeed exclusive eMI-like activity, more evidence on the specificity of these observations is required. First, we investigated whether Venus puncta formation depends on the ESCRT machinery. Hence, siRNA were designed to target RT *tsg101* mRNA, which encodes an essential protein of the ESCRT-I complex^1^. This siRNA (si*tsg101*) efficiently knocked down *tsg101* gene expression in RTH-149 cells (Figure 3A) and reduced its protein levels (Figure 3B; quantification in 3C). Transfection of RTH-149 cells expressing both hemi-KFERQ-Split Venus constructs with negative control siRNA (si*LUC*) did not affect H_2_O_2_-induced puncta formation (Figure 3D; quantification in 3E). However, si*tsg101* prevented the formation of the Venus-puncta. Moreover, siRNAs targeting the two essential and limiting CMA receptors (*lamp2a*-C14 and *lamp2a*-C31), previously validated^20^ and confirmed here (Supplementary Figure 2A, quantification S2B), did not alter Venus puncta formation (Figure 3D; quantification in 3E). Interestingly, blocking CMA with si*lamp2a* significantly (*p*<0.001) increased the number of H_2_O_2_-induced Venus-puncta compared to si*LUC* (Figure 3E), suggesting CMA absence might be compensated by an eMI-like process. This aligns with our prior study, which showed increased Tsg101 levels following *in vitro* silencing of *lamp2a*^20^. Similarly, *tsg101* knockdown increased H_2_O_2_-induced CMA-puncta compared to si*LUC* (Supplementary Figure 2B), suggesting reciprocal compensation between these pathways for degrading KFERQ-containing proteins when one is impaired. This crosstalk between autophagy pathways was also reported in mammals, where Bag6 plays a key role in switching between them^9^. In RT, which has four *bag6* paralogs (Figure 1B), understanding Bag6’s role is a challenge for future research to uncover the molecular mechanisms regulating eMI.

**Figure 3.**
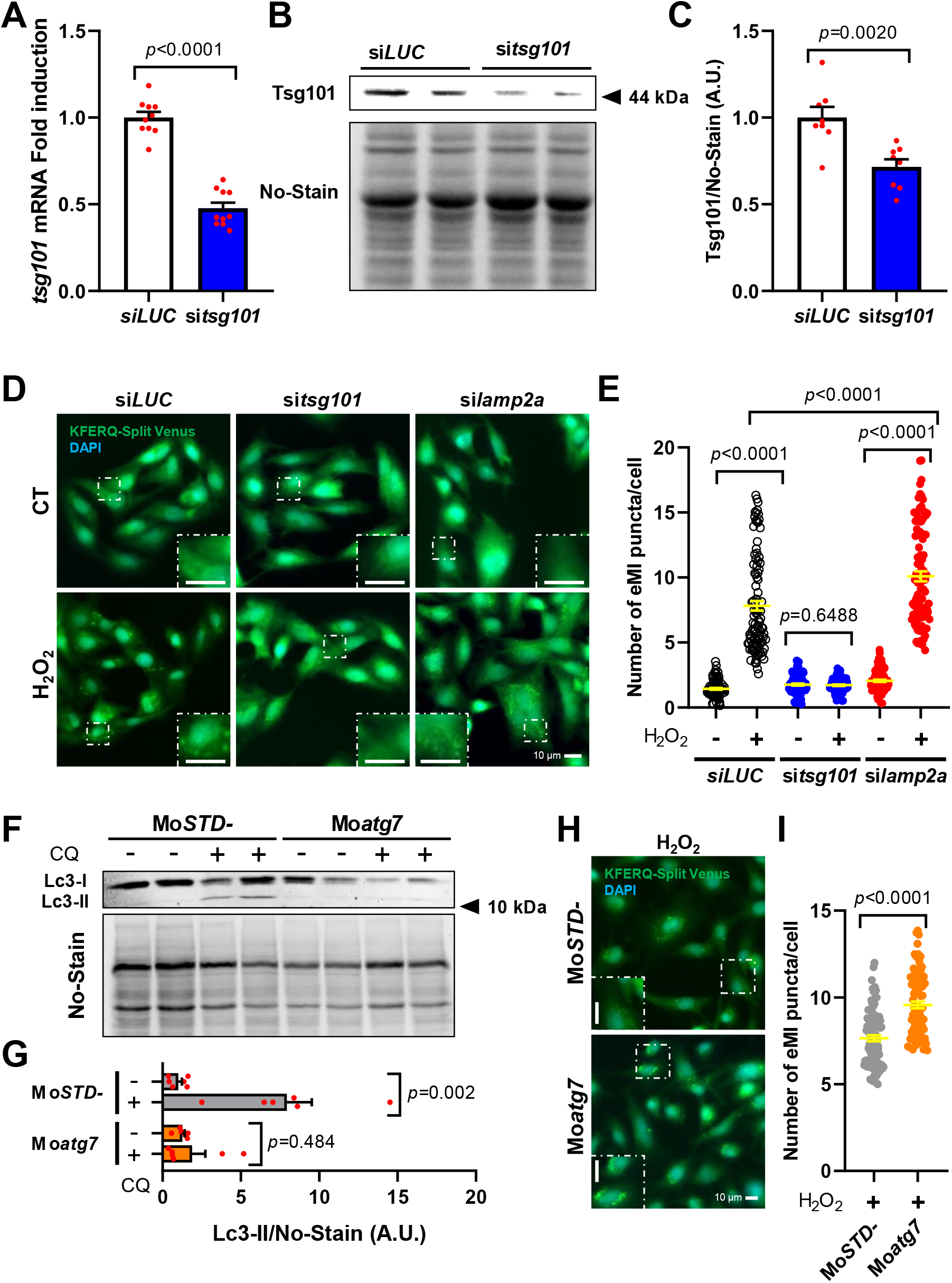
The occurrence of eMI-like puncta under mild-oxidative stress depends on ESCRT machinery, but not on CMA nor on MA. (**A**) The *tsg101* mRNA levels cells were significantly downregulated by the transfection of RTH-149 cells with si*tsg101*. Differences with respect to the negative control condition (si*LUC*) were assessed using Unpaired Student’s T-test (p < 0.0001) in five independent experiments with duplicates. This transfection with si*tsg101* also reduced significantly the proteins levels of Tsg101, as it can be observed in (**B**) the representative western blot image of Tsg101 and the total protein stain (No-Stain), and (**C**) the densitometry quantification of Tsg101 normalized with No-Stain. Differences with respect to the si*LUC* were assessed using Unpaired Student’s T-test (p < 0.002) in four independent experiments with duplicates. (**D**) Representative images of RTH-149 cells expressing the eMI-reporter transfected with either negative control siRNA (si*LUC*), si*tsg101*, or a combination of two siRNAs targeting the two *lamp2a* paralogs (si*lamp2a*) of RT, and incubated with CT medium or exposed to mild-oxidative stress (H_2_O_2_ 25 μM) for 16 h, and the (**E**) quantification of KFERQ-Venus number of puncta per cell. All values correspond to individual images (si*LUC* CT 96; si*LUC* H_2_O_2_ 101; si*tsg101* CT 96; si*tsg101* H_2_O_2_ 96; si*lamp2a* CT 96; si*lamp2a* H_2_O_2_ 95), with >31 images/experiment in a total of 3 independent experiments (> 600 cells per condition). Differences between two groups were assessed using parametric Unpaired Student’s T-test or the non-parametric Mann Whitney test, and the p-values are indicated in the figure. (**F**) Representative image of western blot against LC3B in RTH-149 cells transfected with a morpholino oligonucleotide (Mo) targeting RT Atg7 (Mo*atg7*) or a negative control (Mo*STD-*), exposed during 16 h to H_2_O_2_ 25 μM and additionally incubated with or without 10 μM chloroquine (CQ), and (**G**) quantification of LC3-II protein levels normalized to No-Stain. Differences between groups were assessed using parametric Unpaired Student’s T-test (Mo*atg7* group) or non-parametric Mann Whitney test (Mo*STD-*) in three independent experiments with duplicates. (**H**) Representative images of RTH-149 cells expressing the eMI-reporter transfected with either Mo*STD-* or Mo*atg7* and exposed to H_2_O_2_ 25 μM for 16 h, and (**I**) quantification of KFERQ-Venus number of puncta per cell. All values correspond to individual images (Mo*STD-* 88; Mo*atg7* 90), with >21 images/experiment in a total of 3 independent experiments (> 600 cells per condition). Differences between the two groups were assessed using Mann Whitney test (p < 0.0001). All data are presented as Mean ± SEM; scale bars: 10 μm.

To next evaluate whether the ESCRT-I-dependent but CMA-independent eMI-like puncta might also rely on MA, we designed a morpholino oligonucleotide (Mo) to inhibit the translation of mRNAs encoding the E1-like enzyme Atg7 in the RT. This approach aims to impair MA initiation^39^. We assessed LC3 lipidation as a readout of the MA flux after exposing cells to H_2_O_2_, with or without the autophagosome-lysosome fusion inhibitor chloroquine (CQ, at 10 μM)^40^. Mo*atg7* pointedly reduced LC3-II levels compared to cells transfected with a negative control oligo (Mo*STD-*) (Figure 3F, quantification in 3G), confirming effective MA inhibition. We then examined Venus-puncta formation under mild-oxidative stress and observed that MA inhibition did not decrease the number of eMI-sensor puncta (Figure 3H, quantification in 3I). Instead, it increased the puncta count compared to Mo*STD-*, similar to findings in *Drosophila* fat body^36^. This suggests that increased cellular stress might enhance eMI-like process in RT as well.

Together, our results demonstrate that the formation of the KFERQ-Venus puncta upon H_2_O_2_ incubation specifically relies on ESCRT-I machinery but is indeed independent of both CMA and MA activities, supporting -for the first time-the existence of an eMI-like process in fish. Interestingly, we have observed that the activity of such process in RT increases when either CMA or MA are impaired, which points to an established and probably finely tuned crosstalk between these autophagy pathways to maintain cellular proteome homeostasis. This interaction certainly warrants further research.

### eMI induction in RT is stimulus-specific

As reviewed by Wang et al.^2^, eMI induction appears to be stimulus-specific. In *Drosophila*, eMI is triggered by prolonged nutrient starvation^41^, oxidative stress, or DNA damage^36^. In mammals, eMI can be stimulated by iron starvation and DNA damage^11,37^, but unlike in *Drosophila*, it is either unaffected or downregulated during nutrient deprivation^1,9^. In this study, we examined the effect of different cellular stresses on eMI-sensor puncta formation in RTH-149 cells. We found that high-glucose (HG) treatment, similar to H_2_O_2_, increased the number of Venus puncta compared to the CT condition (Figure 4A, quantification in 4B). We recently showed that HG induces oxidative stress in these cells, mirroring H_2_O_2_ effects, and reflecting the glucose intolerance typical of RT^20^. Notably, existing data from *Drosophila* suggest a MAPK/JNK signaling involvement in eMI induction during oxidative stress^36^. However, the potential role of NRF2 cannot be ruled out, as demonstrated for CMA in mammalian cells^42^, and more recently in RTH-149^20^. Future research should explore if this mechanism is conserved in RT.

**Figure 4.**
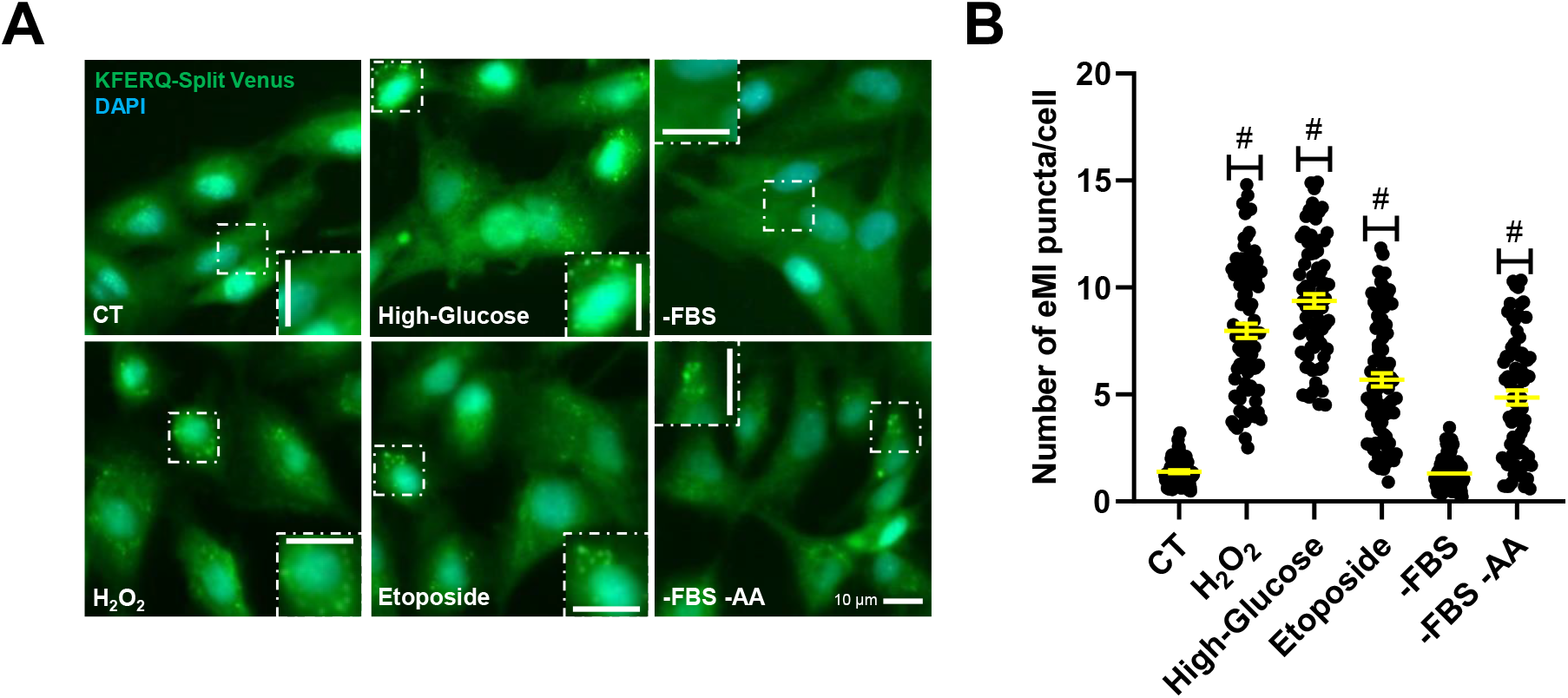
Different cellular stressors, although not serum starvation, activated eMI-sensor puncta formation in RTH-149 cells. (**A**) Representative images of RTH-149 cells expressing the eMI-reporter and incubated for 16 h with either CT medium, H_2_O_2_ (25 μM), glucose (25 mM, High-Glucose), etoposide (50 μM), a medium lacking serum (-FBS) alone, or a medium lacking both serum and amino acids (-FBS –AA). (**B**) Quantification of KFERQ-Venus number of puncta per cell. All values correspond to individual images (CT 80; H_2_O_2_ 86; High-Glucose 76; Etoposide 86; –FBS 87; –FBS –AA 76), with >15 images/experiment in a total of 3 independent experiments (> 600 cells per condition). Differences between two groups were assessed using the non-parametric Mann Whitney test, and the p-values are indicated in the figure. All data are presented as Mean ± SEM; scale bars: 10 μm.

We then examined the effect of genotoxic insults by inducing DNA double-strand breaks with etoposide (50 μM)^43^. Our results showed that etoposide clearly increased Venus puncta compared to CT (Figure 4A, quantification in 4B), similarly to findings in *Drosophila*^36^ and mammals^37^. In mammals, CMA participates in the selective degradation of the checkpoint kinase 1 (Chk1), a KFERQ-motif containing protein, to ensure cell cycle progression after DNA repair^44^. Conversely, CMA impairment was associated with defects in nuclear proteostasis, genomic instability and reduced cell survival^44^. Although it remains to be determined whether Chk11 can also be targeted by eMI, these findings suggest a conserved protective mechanism that maintains genome integrity through the degradation of KFERQ-substrates after DNA damage, which could be, at least partly, fulfilled by eMI in RT.

Finally, we observed that serum starvation (-FBS), known to induce both CMA^3,45,46^ and MA in various species, including RT^47–49^, did not alter Venus-puncta formation. In contrast, combined serum and amino acids deprivation (-FBS -AA) with HBSS medium for 16 h substantially increased eMI-like puncta compared to CT (Figure 4A, quantification in 4B). This aligns with increased eMI during prolonged nutrient deprivation in the CMA-deficient species *Drosophila*^41^. In mammals, where both CMA and eMI pathways are present, the regulation of eMI during nutrient deprivation is more complex^28^. Short-time (1 h) HBSS exposure triggers an ESCRT-III-dependent, but ESCRT-I and Hsc70-independent, eMI variant in human cell lines. This variant likely degrades selective autophagy receptors (e.g., LC3B and GABARAPL2), preventing unwarranted selective MA in the short-term, while enhancing bulk MA during prolonged starvation^50^. Conversely, long-term nutrient starvation, a well-known CMA activator^31^, inhibits eMI in rodents, likely due to the CMA-mediated degradation of the eMI component BAG6^9^. This research highlights the possible role of BAG6 as a switcher, directing KFERQ-containing proteins to either eMI or CMA and enabling these pathways to compensate for each other’s deficiencies. In RT, both autophagic pathways function simultaneously during stress, such as mild-oxidative stress and nutrient deprivation (16 h), targeting KFERQ-containing proteins to lysosomes and LE/MVB. This raises the question of whether a Bag6-dependent mechanism coordinates eMI and CMA pathways in RT as observed in mammals. However, the differences between our results and those in mammals may stem from the experimental models used. In our *in vitro* model, -FBS -AA represents an acute condition with negligible extracellular levels of these compounds, while mammalian studies involved *in vivo* nutrient starvation experiments^9^, where circulating nutrient and growth factor levels likely remain physiological. To the best of our knowledge, no data exists on the *in vitro* activity of eMI in mammals during both long-term and acute nutrient deprivation. Therefore, we cannot been ruled out that the simultaneous activation of CMA and eMI observed in the present study may be a conserved physiological response regarding extreme cellular stress across vertebrates.

In summary, we identified and characterized an eMI-like pathway in a RT hepatocyte cell line. This process is induced by oxidative stress, DNA damage, or nutrient deprivation, mirroring findings in *Drosophila*^36,41^, but not in response to FBS removal, supporting that eMI induction in RT is stimulus-specific. These findings suggest a significant role for this eMI-like process in fine-tuning the cellular proteome during stress in RT. Moreover, they open new avenues for exploring this function using complementary model organisms, such as zebrafish and RT. Further studies are needed to determine whether this process depends on KFERQ motifs and HSC70 and to investigate the potential role of the RT Bag6 paralogs.

## Materials and methods

### Laboratory ware and reagents

Minimum Essential Medium (61100-053), essential (11130-036) and non-essential amino acids (11140-050), L-glutamine (25030-024), sodium pyruvate (11360-070), penicillin-streptomycin (15140-122), geneticin (11811031), fetal bovine serum (10270-106), and Hank’s Balanced Salt Solution (14025-092), were all purchased from Gibco. HEPES (BP299-1) and PBS (BP2944) were procured by Fisher BioReagents. Hydrogen peroxide solution (H1009), D-(+)-Glucose (G7021) and chloroquine (C6628) were all purchased from Sigma-Aldrich. Etoposide (HY-13629) was purchased from MedChemExpress, paraformaldehyde 4% fixative (22023) from Biotium, and the antifade mounting medium containing DAPI (H-2000) from Vector Laboratories, Inc. The TRIzol Reagent (15596018), RIPA buffer (89901), the cocktail of protease and phosphatases inhibitors (78422), the Qubit protein assay kit (Q33211), SuperBlock Blocking Buffer in PBS (37515), SuperSignal West Pico Plus Chemiluminescent Substrate (34578), and No-Stain Protein Labeling Reagent (A44449), were all acquired from ThermoFisher Scientific. All laboratory plastic ware, unless otherwise stated, were procured by Sarstedt AG & Co. KG.

### Bioinformatics analysis, oligos and plasmids

Ensembl Genome database (Ensembl release 112; http://www.ensembl.org) was used to identify orthologous genes to the human genes coding for the main eMI players in the genome of RT (Genome assembly: USDA_OmykA_1.1), or of *Danio rerio* (Genome assembly: GRCz11), and their identity was later confirmed using Genomicus version 110.01^51^. To screen whether the identified genes are expressed across different tissues of RT and zebrafish, we interrogated the PhyloFish RNA-seq database^35^. Instead, to evaluate their expression in RTH-149 cells, we performed a Bulk RNA barcoding and sequencing (BRB-seq) analysis^52^ using the services of Alithea Genomics S.A as explained below.

Regarding oligos, up to four new siRNAs against the RT *tsg101* mRNA sequence (Ensembl ID-gene: ENSOMYG00000006944) were designed using the siDESIGN Center. After testing their efficiency in reducing both mRNA and protein levels of Tsg101 by qPCR and western blot (data not shown), the most effective was selected for following experiments (si*tsg101*; 5’-UGAUUAGGUGAGCGAAUUA-3’). The different siRNAs using in the present work, including the siRNAs against the two RT *lamp2a*’s (i.e., si*14*; 5’-UAAAGAAAUUGCUCGGCUC-3’, and si*31*; 5’-UGGAGAAGCGGCUGUGUUA-3’, previously validated for RT^20^, and a negative control siRNA against LucGL2 (si*LUC*; 5’-CGUACGCGGAAUACUUCGA-3’), not expressed in RT cells, were purchased from Horizon Discovery. Moreover, a morpholino oligonucleotide (Mo*atg7*; 5’-CAGAAGCCATCTCTATCACCG-3’) was designed in collaboration with GeneTools customer support to targeting the start codon of the RT *atg7* mRNA sequence (Ensembl ID-gene: ENSOMYG00000075941) for blocking its translation. This oligo and a standard negative control (Mo*STD-*; 5’-CCTCTTACCTCAGTTACAATTTATA-3’) targeting an intron of human beta-globin associated with beta-thalassemia were purchased from GeneTools.

Concerning plasmids, two constructions originally designed for tracking eMI in *Drosophila*^33^ and based on the backbone pUAST-attB (RRID:DGRC_1419) plus the 21^st^ amino acids of RNASe A, including its KFERQ motif, fused to either the N-or C-Split Venus^53^, were gifted by Patrik Verstreken (KU Leuven). Those two plasmids were digested to isolate the N- and C-Split Venus regions tagged to the KFERQ motif, for their posterior sub-cloning in the backbone of the pEGFP-N1-ro1GFP [a gift from S. James Remington (Addgene plasmid #82369; http://n2t.net/addgene:82369; RRID:Addgene_82369)] for a suitable expression in RT cells, but only after the removal of the EGFP sequence. Moreover, mRFP-Rab7 plasmid was a gift from Ari Helenius (Addgene plasmid #14436; http://n2t.net/addgene:14436; RRID:Addgene_14436).

### Cells and general experimental procedures

The RT hepatoma-149 cell line (RTH-149 cells; supplied by ATCC #CRL-1710; RRID:CVCL_3467), passage number below 20, and routinely screened to confirm absence of mycoplasma contamination by using the Myco-Sniff Mycoplasma PCR Detection Kit (MP Biomedicals #093050201) following the manufacturer’s recommendations, was used for all cell culture experiments in the current work. This cell line has been previously validated as a reliable model for nutrition and autophagy research in the RT^20,48^. Cells were grown at 18°C without CO_2_ in complete medium (hereafter referred as control condition, CT) consisting in Minimum Essential Medium containing essential amino acids, 1X L-glutamine and glucose (5 mM), and supplemented with 1X non-essential amino acids, sodium pyruvate (1 mM), HEPES (25 mM), 1X Penicillin-Streptomycin, and 10% fetal bovine serum.

RTH-149 cells stably transfected with the two N- and C-Split Venus regions tagged to the KFERQ motif were selected for the resistance to the selective antibiotic geneticin, followed by cell sorting using a FACS Aria 2-Blue 6-Violet 3-Red 5-YelGr, 2 UV laser configuration (BD Biosciences, Le Pont de Claix Cedex, France) in biosafety cabinet. On the other hand, RTH-149 cells stably expressing a KFERQ-PA-mCherry1 generated and selected in our previous work^20^ were additionally used for tracking CMA. Those eMI-or CMA-reporter cells, respectively, were maintained in a version of the CT medium detailed above in which the antibiotic used was geneticin. All the cells transfections with either plasmids (1-2.5 μg), siRNAs (1 μM) or morpholino oligos (5 μM), were performed with the Transfection System device (Invitrogen MPK5000) and Neon 100 μL samples Kit (Invitrogen MPK10096).

At the beginning of each experiment, similar to what was previously detailed in Vélez et al.^20^, cells were counted using a Cellometer K2 device (Bioscience LLC, Lawrence, MA, USA) and plated at a density of 60.000 cells/well onto 4-well culture slides (Ibidi 80426 or FALCON 354114) for Venus or mCherry microscope imaging experiments, respectively. In the case of RNA extraction and protein collection experiments, cells were plated in 6-cm dishes at 400.000 or 500.000 cells/dish, respectively. Before experimental treatments, the cells were gently washed twice with 1X PBS, and then incubated with CT medium alone, or supplemented with either hydrogen peroxide (25 μM, H_2_O_2_ medium, mild-oxidative stress), High-Glucose (D-glucose at 25 mM), or etoposide (50 μM), as appropriate and for the times detailed elsewhere. Otherwise, cells were incubated with H_2_O_2_ medium supplemented with chloroquine (CQ condition, 10 μM), or instead, with 1X Hank’s Balanced Salt Solution supplemented exclusively with HEPES (25 mM, HBSS condition, serum and nutrient starvation medium), or also with 1X essential amino acids, 1X non-essential amino acids, and 1X L-glutamine (-FBS condition, serum starvation medium). To asses CMA activity, the CMA-reporter expressing cells were photoactivated with a 405-nm light source for 10 min prior to starting the incubations^20^.

### Imaging procedures

Cells were fixed for 20 min using paraformaldehyde 4% before mounting and counterstaining nuclei with DAPI, and then the images were acquired in the Bordeaux Imaging Center (CNRS-INSERM and Bordeaux University, member of the national infrastructure France BioImaging). On the one hand, all cells images of KFERQ-Venus and mRFP-Rab7 fluorescence were acquired in widefield using a 20X/0.95 dry objective and a 1X intermediate lens with the high-content screening microscope Zeiss Cell Discoverer 7 (Zeiss Microscopy, Germany), which is equipped with Dual Zeiss Colibri LED-based illumination (from 385 nm to 625 nm), the dichroic mirrors 405/493/575/653, 450/538/610, 405/493/610, the emission filters 412-438/501-527/583-601/662-756, 458-474/546-564/618-756, 412-433/501-547/617-758, the color camera Zeiss Axiocam 512 as a detector, and its controlled by the Zen Blue 3.1 software (Zeiss). On the other hand, as reported before20, for imaging CMA-reporter cells we used a widefield Microscope Leica DM 5000 (Leica Microsystems, Germany) equipped with a 40X/1.25 oil objective, DAPI and TRITC filter cubes, a Greyscale sCMOS camera (Hamamatsu Flash 4.0 V2, Japan), and controlled by MetaMorph v7.8.10.0 software (RRID:SCR_002368).

eMI-reporter puncta per image were automatically detected in the Venus-channel and counted with Fiji (RRID:SCR_002285) by using the ComDet v.0.5.5 plugin (developed in Cell Biology group of Utrecht University and available at https://github.com/UU-cellbiology/ComDet), with an approximate particle size of 4 pixels and an intensity threshold ranging from 5 to 20 SD as settings. Next, the number of nucleus (> 100 pixels) in each image were counted by using the Fiji Analyze Particles function in the DAPI channel after IsoData thresholding and watershed segmentation. In the case of PA-mCherry1 fluorescence experiments, the CMA puncta were counted on images in gray (red channel) and blue (DAPI) by using the Cell Counter plugin of Fiji (RRID:SCR_002285) as previously detailed20, followed by nuclei counting. All images were counted in blind and random order and excluding cells on the edges or very superposed. For each image, the number of either eMI-or CMA-puncta was normalized by the number of nuclei within the image, and the results showed represent the number of puncta per cell in a minimum of 60 images coming from 3 independent experiments. In all cases, more than 600 cells/condition were counted.

The BIOP version (https://github.com/BIOP/ijp-jacop-b) of the original JACoP plugin (RRID:SCR_025164) for Fiji was used to evaluate the co-ocurrence between Venus-puncta and late endosomes labeled with mRFP-Rab7 in cells incubated with H2O2 medium. We analyzed in 85 single cells the PCC, which ranges from −1 to 1, where −1 indicates a negative correlation and +1 highlights a complete positive correlation, between the pixel-intensity of the two contrasted channels38.

### mRNA levels analysis: BRB-seq and quantitative RT-PCR

RNA samples were collected and processed with TRIzol reagent following the manufacturer’s protocol. Then, RNA concentration was determined using a NanoDrop 2000 (Thermo Scientific) and integrity confirmed in an agarose gel, as previously reported^54^. In the case of Bulk RNA barcoding and sequencing (BRB-seq) analysis^52^, 8 independent RNA samples (50 ng of RNA/each) were collected and isolated from RTH-149 cells cultured in growth medium and successively sent to Alithea Genomics S.A for its processing as explained in detail elsewhere^22^. On the other hand, for quantitative RT-PCR the samples were prepared as described before^48,55^ and cDNA synthesis was performed in duplicate, followed by duplicated PCR analysis^20^. The relative quantification of the target gene *tsg101* was calculated using the ΔCT method^56^ with *eef1a1* (eukaryotic translation elongation factor 1 alpha 1) as a reference gene. Primers used for *tsg101* (forward: 5’-CCAGTCAGCTACAGAGGAAACA-3’ / reverse: 5’-AGATCTTGCCGTTGGCATCA-3’) were designed in the current work by using the Primer-Blast tool of NCBI, and the amplicon was sent to Genewiz (Azenta Life Sciences) to confirm primer specificity by Sanger Sequencing. Primers used for *eef1a1* (forward: 5’-TCCTCTTGGTCGTTTCGCTG-3’ / reverse: 5’-ACCCGAGGGACATCCTGTG-3’) were previously validated^57^.

### Western blotting

Proteins were collected with RIPA buffer supplemented with protease and phosphatases inhibitor^48^, and then quantified using a Qubit 2.0 fluorometer (ThermoFisher Scientific). Later, 5 μg of protein were separated by sodium dodecyl sulfate polyacrylamide gel electrophoresis (SDS-PAGE) and then proteins were transferred to polyvinylidene fluoride (PVDF) membranes (Merk-Millipore IPFL00010). Next, total protein labeling was performed with No-Stain Protein Labeling Reagent following the manufacturer’s protocol for posterior accurate total protein normalization. After blocking membranes with SuperBlock Blocking Buffer in PBS, they were incubated with the corresponding primary antibody rabbit anti-TSG101 (Sigma-Aldrich HPA006161, at 1/500), or rabbit anti-LC3B (Cell Signaling Technologies 2775, at 1/1,000), followed by incubation with a goat anti-rabbit secondary antibody HRP (Invitrogen 31460, at 1/10,000). Immunoreactive bands were developed thanks to the SuperSignal West Pico Plus Chemiluminescent Substrate, and imaged using Smart Exposure to assure signal linearity in an iBright FL1500 Imaging System (ThermoFisher Scientific). Finally, the images were quantified using the iBright Analysis Software (ThermoFisher Scientific, RRID:SCR_017632).

### Statistics and Reproducibility

All statistical analyses in the current study were performed using GraphPad Prism (RRID:SCR_002798) version 8.0.1 for Windows (GraphPad Software, Inc., www.graphpad.com) considering a *p*-value < 0.05 as a level of significance, and all data presented are reported as means ± SEM or fold induction respect the control group ± SEM, from a minimum of 3 independent experiments. Before performing parametric or non-parametric tests, Normal distribution was assessed. The differences between two groups were evaluated through a parametric two-tailed unpaired Student’s T-test, or a non-parametric Mann Whitney test. Instead, for the comparison between more than two groups in Figure 2B, Kruskal-Wallis followed by Dunn’s multiple comparisons test was utilized.

## Data Availability

The datasets generated during and/or analyzed during the current study are openly available, along with their statistic report, in the Recherche Data Gouv repository, at 58 https://doi.org/10.57745/K4XF9I.

## Acknowledgments

This project has received funding from the European Union’s Horizon 2020 research and innovation programme under the Marie Sklodowska-Curie grant agreement No 101030643, and Aquaexcel3.0 grant agreement No 871108. We are thankful to P. Verstreken for the original Venus plasmids, to S.J. Remington for the EGFP-N1-ro1GFP, and A. Helenius for the mRFP-Rab7. We thanks V. Pitard [Flow cytometry facility], as well as C. Poujol and F. Cordelières [Bordeaux Imaging Center (BIC), France BioImaging national infrastructure member supported by the French National Research Agency (ANR-10-INBS-04)], from CNRS-INSERM, University of Bordeaux. We thanks L. Peron for the light emitting device for Pa-mCherry photoactivation, and J. Bugeon (INRAE LPGP) for helping us with the automation of fluorescent puncta count. Schematic Figure 1A was created with BioRender.com tools.

## Abbreviations

CMA: chaperone-mediated autophagy
CQ: chloroquine
eMI: endosomal microautophagy
HG: high-glucose
LAMP2A: lysosomal associated membrane protein 2A
LE: late endosome
MA: macroautophagy
Mo: morpholino oligonucleotide
MVB: multivesicular bodies
PCC: Pearson’s correlation coefficient
RAB7: Ras-related protein 7
RT: *Oncorhynchus mykiss*

## Author contributions

EJV and IS conceived the study, obtained the funding and prepared the first draft. EJV, VV and JG performed the *in vitro* experiments and laboratory analyses. All authors (EJV, VV, JG, SR, KD, AH, FB, and IS) contributed to the discussion and edition, and approved the final manuscript.

## Supplementary information

Supplementary Figure 1 shows that the core eMI machinery is present in the zebrafish genome and expressed in different tissues. Supplementary Figure 2 shows the effects of *tsg101* or *lamp2a* siRNA-mediated silencing on the formation of CMA puncta in RTH-149 cells.

**Supplementary Figure 1.**
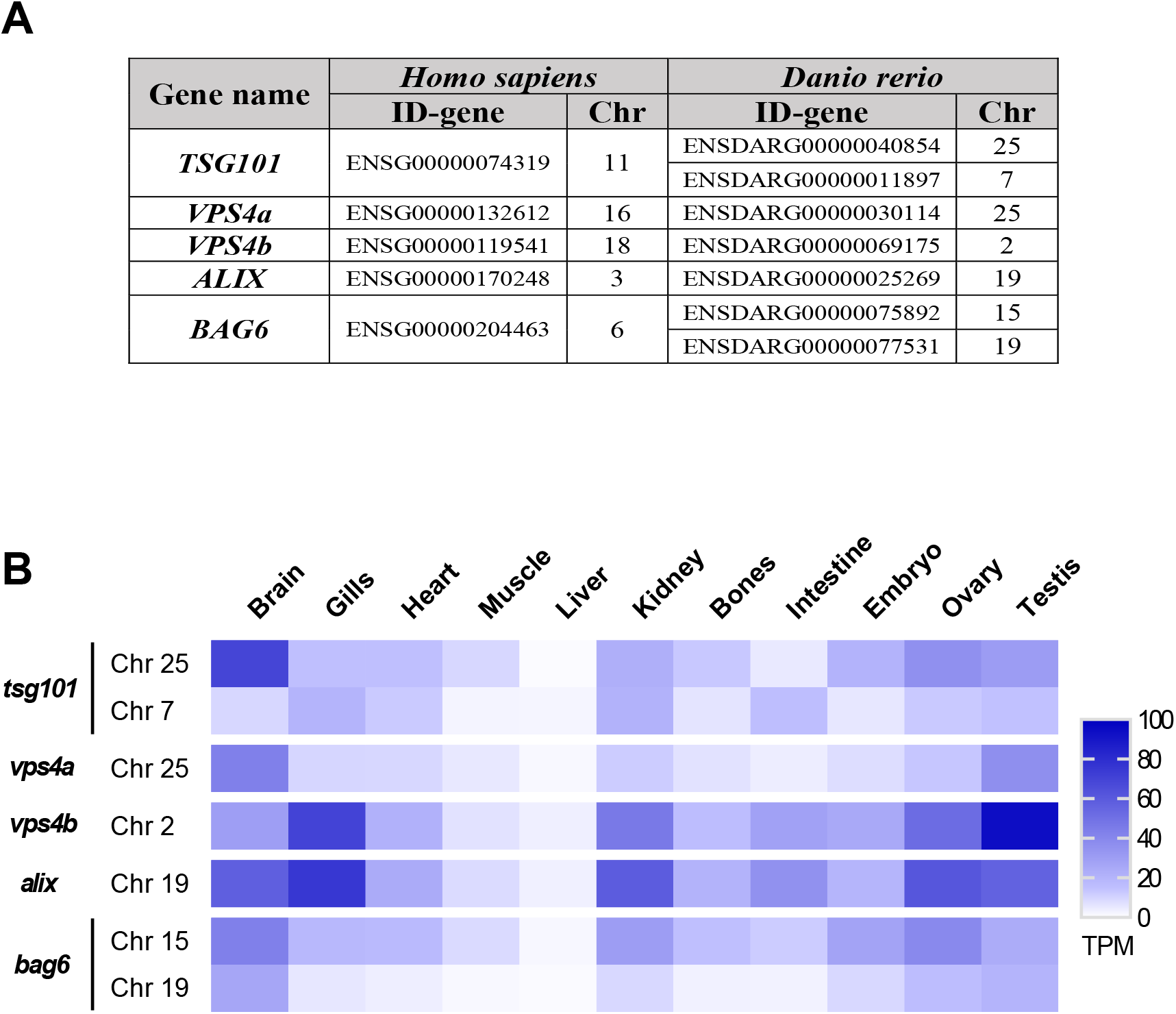
The core eMI machinery is present and expressed in *Danio rerio* (zebrafish) tissues. (**A**) Identification (Ensembl ID) and chromosome location (Chr) of the main eMI-genes (*TSG101, VPS4A, VPS4B, ALIX* and *BAG6*) in the human genome, and their corresponding zebrafish orthologs. (**B**) Heat map showing mRNA expression levels (TPM, transcripts per million) of the different identified genes in different zebrafish tissues (brain, gills, heart, muscle, liver, kidney, bones, intestine, embryo, ovary and testis) extracted from the RNA-Seq database PhyloFish.

**Supplementary Figure 2.**
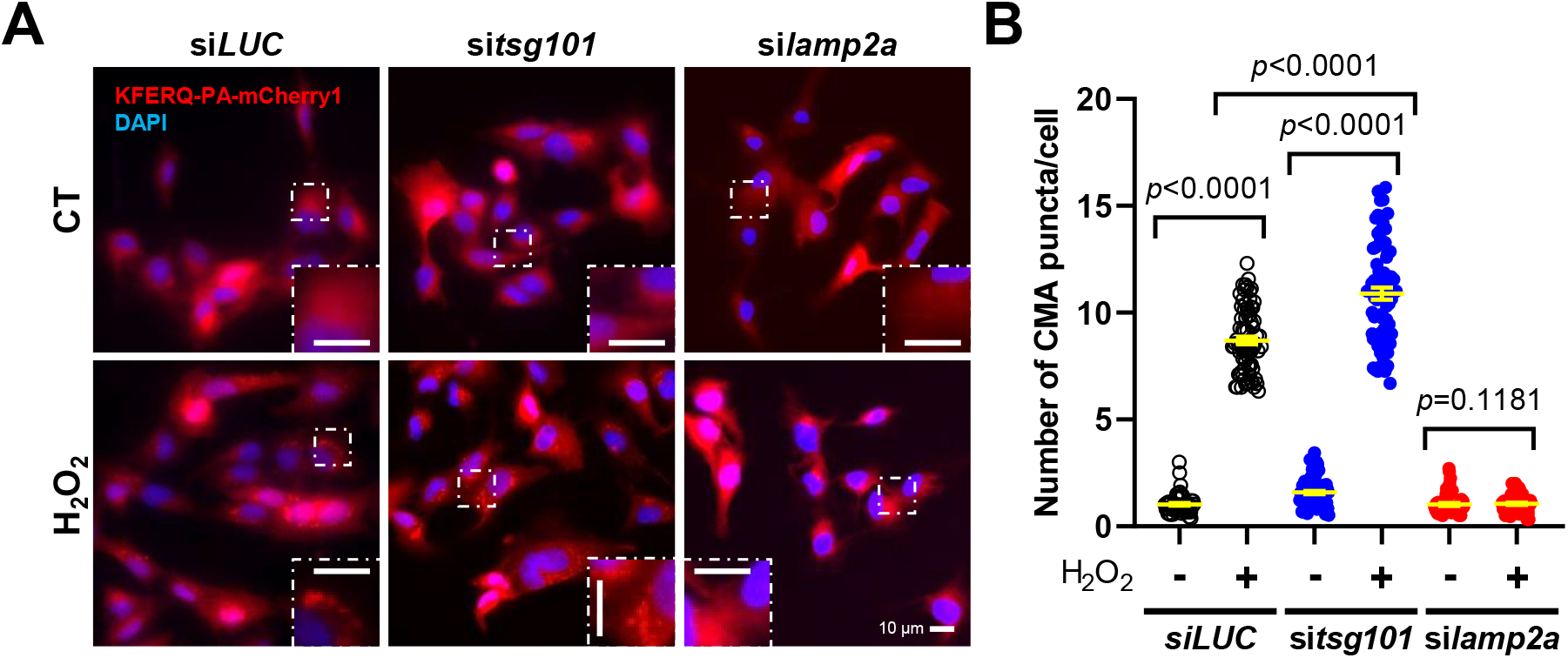
The silencing of the ESCRT-I protein Tsg101 by using si*tsg101* did not prevented the formation of CMA puncta induced by mild-oxidative stress exposure. (**A**) Representative images of RTH-149 cells expressing the KFERQ-PA-mCherry1 CMA-reporter and transfected with either negative control siRNA (si*LUC*), si*tsg101*, or a combination of two siRNAs targeting the two *lamp2a* paralogs (si*lamp2a*) of RT, and incubated with CT medium or exposed to mild-oxidative stress (H_2_O_2_ 25 μM) for 16 h, and the (**B**) quantification of CMA number of puncta per cell. All values correspond to individual images (si*LUC* CT 60; si*LUC* H_2_O_2_ 77; si*tsg101* CT 65; si*tsg101* H_2_O_2_ 70; si*lamp2a* CT 60; si*lamp2a* H_2_O_2_ 55), with >20 images/experiment in a total of 3 independent experiments (> 600 cells per condition). Differences between two groups were assessed using the non-parametric Mann Whitney test, and the p-values are indicated in the figure. All data are presented as Mean ± SEM; scale bars: 10 μm.

